# Self-Contemplating In-Context Learning Enhances T Cell Receptor Generation for Novel Epitopes

**DOI:** 10.1101/2025.01.27.634873

**Authors:** Pengfei Zhang, Seojin Bang, Heewook Lee

## Abstract

Computational design of T cell receptors (TCRs) that bind to epitopes holds the potential to revolutionize targeted immunotherapy. However, computational design of TCRs for novel epitopes is challenging due to the scarcity of training data, and the absence of known cognate TCRs for novel epitopes. In this study, we aim to generate high-quality cognate TCRs particularly for *novel epitopes* with no known cognate TCRs, a problem that remains under-explored in the field. We propose to incorporate in-context learning, successfully used with large language models to perform new generative tasks, to the task of TCR generation for novel epitopes. By providing cognate TCRs as additional context, we enhance the model’s ability to generate high-quality TCRs for novel epitopes. We first unlock the power of in-context learning by training a model to generate new TCRs based on both a target epitope and a small set of its cognate TCRs, so-called in-context training (ICT). We then self-generate its own TCR contexts based on a target epitope, as novel epitopes lack known binding TCRs, and use it as an inference prompt, referred to as self-contemplation prompting (SCP). Our experiments first demonstrate that aligning training and inference distribution by ICT is critical for effectively leveraging context TCRs. Subsequently, we show that providing context TCRs significantly improves TCR generation for novel epitopes. Furthermore, we show TCR generation using SCP-synthesized context TCRs achieves performance comparable to, and sometimes surpassing, ground-truth context TCRs, especially when combined with refined prompt selection based on binding affinity and authenticity metrics.

## 1. Introduction

Genetically engineered T cells equipped with therapeutic T cell receptors (TCRs) have emerged as a transformative approach in personalized immunotherapy for treating diseases such as cancer [2, 21]. Cognate TCRs play crucial roles for T cells in the identification of abnormal cells by recognizing disease-specific epitopes–an epitope is a part of antigen that sits in the binding pocket of cognate receptor– presented by major histocompatibility complex (MHC) on cell surface [1] (Figure 1A). Computational generation and validation of cognate TCRs for target antigens can expedite the process of developing personalized engineered T cells (Figure 1B). It significantly reduces the number of candidate TCRs subject to wet-lab validation, resulting in substantial reductions in time and cost. It particularly presents a unique opportunity for *novel epitopes*, where timely identification of cognate TCR is essential. Novel epitopes, by definition, are peptides with no known cognate TCRs. Examples include those from newly emerging pathogenic viral strains or patient-specific cancer-induced neoantigens.

**Fig. 1.**
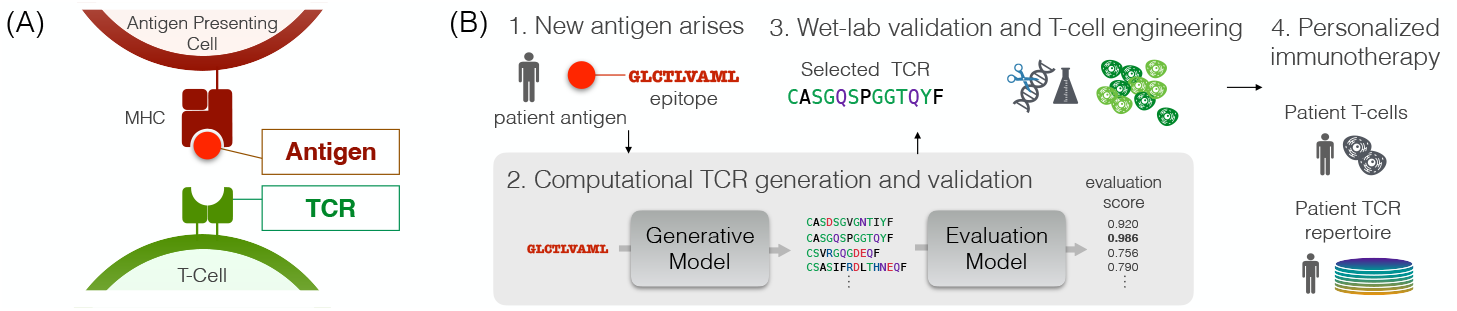
(A) TCR, a protein complex located on the surface of T cells, recognizes and binds to a specific part of antigen (epitope). It allows T-cell to recognize and kill abnormal cells. (B) This study investigates computational generation and validation of TCRs (gray background) within personalized immunotherapy development. Personalized immunotherapy development is powered by the computational approaches: The computational approaches prior to wet-lab validation can offer substantial cost savings and improved workflow efficiency.

Cognate TCR generation requires reliable, rapid validation of generated TCRs. Recent success in predicting the binding affinity of TCR epitopes [11, 17, 4, 24, 23, 25], with a recent work achieving high accuracy (AUC *>* 0.94 even for novel epitopes), opens a door to tackling the generation task. Despite promising advances in the affinity validation, the generation of cognate TCRs remains largely unexplored. Current efforts are limited to generation of TCR repertoires (not considering cognate epitopes), or only in cases where known TCR-epitope pairs exist [6, 8].

TCR generation for novel epitopes requires models with enhanced generalization capacity in order to produce TCRs fore previously unseen and potentially out-of-distribution epitopes. Large language models (LLMs) such as GPT, demonstrated remarkable success in generalizing to new tasks, stemming from their training on extensive and diverse datasets [3]. LLMs adapt to new tasks via in-context learning, dynamically adjusting responses on the context of a new task, typically provided as a few examples of the task during inference [3]. However, applying LLMs to TCR generation for novel epitopes presents unique challenges. Existing datasets of TCR-epitope pairs cover only a limited range of epitopes and each epitope has far fewer corresponding TCRs than it can recognize. The scarcity of data hinders generalization to novel epitopes which are unseen during training. Furthermore, the lack of known TCRs precludes the use of standard in-context learning for novel epitopes, since it relies on providing contextual TCR examples during inference.

Our goal is to develop a computational model that can generate cognate TCR sequences for novel epitopes. The core idea is to provide the model with both the target epitope sequence and a few of its associated TCRs as input. This enables the model to leverage additional information from the provided TCR context when generating cognate TCRs for the target epitope.

We address two challenges encountered in generating TCRs for novel epitopes. First, to overcome the data scarcity and enhance generalizability to unseen epitopes, we align the model’s training with our inference goal. It involves training a model to generate a new TCR based not only on a target epitope, but also a small set of its cognate TCRs as additional input context, and so-called called in-context training (ICT)^1^. Next, to address the issue of unavailability of known context TCRs for novel epitopes, we eliminate the need for having known binding TCRs at the inference time by leveraging self-contemplation prompting (SCP). It begins self-generating an initial set of TCRs based only on a given epitope, from which high-quality TCRs are selected and used as input context (prompt). We assess the generated TCRs’ authenticity and binding potential using state-of-the-art computational methods as a proxy for wet-lab validation.

This study pioneers the generation of potentially cognate TCRs for novel epitopes, a challenging task because these epitopes fall outside the distribution of known epitopes. We believe our work establishes a foundation for developing more sophistracted TCR generative models for novel epitopes. Further research is needed to refine and validate generated TCRs. Crucially, wet-lab validation remains essential to confirm the functionality of generated TCRs. Code and models are available at a public repository^2^.

## 2. Related Works

We first discuss recent efforts and challenges in computational generation of TCRs. Next, we briefly discuss strengths and limitations of in-context learning and explicit meta-learning to enhance generalizability

### Computational generation of cognate TCRs

While generative models have been widely applied to the generation of biological sequences, including protein sequences [9, 10], the specific domain of the generation of TCR sequences has not been thoroughly explored. Existing models primarily focus on replicating the distribution of real TCR sequences, often overlooking the critical need for epitope specificity [8]. Although reinforcement learning methods have been developed to generate epitope-specific TCRs, their application is restricted to in-sample epitopes, and they require retraining for each new epitope [6]. Importantly, there is a lack of research on computationally generating TCRs for out-of-distribution epitopes, despite the significant interest in TCR engineering for newly emerging epitopes.

The cost of wet-lab validation is another significant challenge in the development of TCR generation models. To address this, computational methods for predicting binding affinity have emerged as valuable proxies. Although not infallible, they expedite the development process by reducing the number of candidates requiring subsequent wet-lab validation. Various computational approaches have been proposed [4, 11, 17]. Recent works, such as catELMo embeddings with shallow linearly connected layers [23] and PiTE [24], which leverage pre-trained TCR embedding models, have achieved an average AUC score of over 94% for TCR-epitope binding prediction, even for unseen epitopes.

### In-context learning

LLMs have been recognized task-agnostic few-shot learners [3]. They can infer outputs for test inputs based on a prompt consists of a few input-output pairs demonstrating a desired task. It particularly beneficial for novel tasks where fine-tuning data is scarce or unavailable. This capability is largely attributed to their extensive pre-training on diverse and large datasets, allowing them to meta-learn implicitly and generalize quickly from few examples [3]. However, when a model is not sufficiently scaled or applied to domains significantly different from their pre-training data, their generalization ability diminishes [26].

Explicit meta-learning, as opposed to the implicit methods, involves pre-training or fine-tuning the model on training data closely aligned with the testing data. Although it requires adjustments of the pre-training parameters, they can significantly improve performance on targeted tasks. A recent work [14] leverages meta-learning to train an LLM for few-shot learning tasks, which they called *meta-training for in-context learning*. Another team [5] proposed a similar approach, which they called *in-context tuning*, involves fine-tuning of LLMs on a small dataset of input-output pairs, enhancing its few-shot inference capabilities.

## 3. Data

We collect experimentally validated TCR-epitope pairs from three publicly available databases: McPAS [19], IEDB [20], VDJdb [16]. We exclude pairs of which epitopes are associated with fewer than 100 TCRs. It results in the removal of 836 unique epitopes, which is 6.89% of the total pairs (Figure 2). We further preprocess the data as described in [4]. Pairs of human MHC class I epitopes (linear) and TCR*β* sequences are included only. Pairs containing wildcards such as `*` or `X` and VDJdb pairs with zero confidence scores are excluded. After preprocessing, our dataset comprises about 140K unique TCR-epitope interactions. To assess how well our models can generalize to *novel epitopes*, we divide the 146 unique epitopes into three distinct sets: training (64%), validation (16%), and test (20%). Each set contains a completely distinct list of epitopes and their associated TCRs. This ensures the models are trained on a representative subset of data and tested on entirely *unseen epitopes*–epitopes that do not appear in training. This results in about 96.7K training, 23.5K validation, and 19.7K test TCR-epitope pairs. We also source four million TCR sequences (unlabeled) from healthy human TCR repertoire (ImmunoSeq, [15]) for developing our evaluation framework. Throughout this paper, we use the term ‘TCRs’ to refer to TCR *β* CDR3 (complementarity-determining region 3) sequences as it is the most important CDR in antigen recognition.

**Fig. 2.**
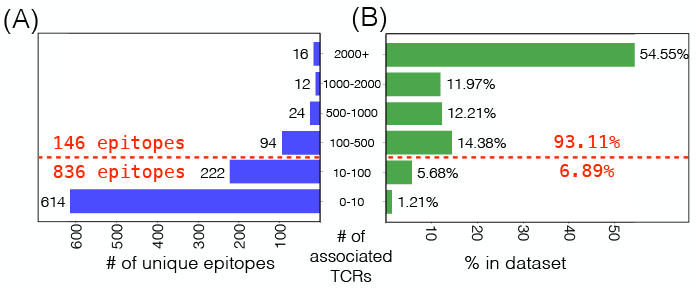
(A) The number of unique epitopes plotted against the number TCRs that recognize them and (B) percentage of each category in the dataset.

## 4. Method

Our approach is in twofold: (1) in-context training to maximize generalizability of in-context inference for TCR generation in Section 4.1 and (2) self-contemplation prompting to enable in-context inference even for novel epitopes without known cognate TCRs in Section 4.2. We then outline our evaluation metrics and baselines to assess the quality of the generated TCRs in Section 4.3.

### 4.1. In-context training

Due to the limited diversity of the training epitopes, the model has difficulty leveraging cognate TCRs provided as in-context information when generating TCRs for unseen epitopes. We address this challenge by training our model in the way that mirrors the inference process itself. This approach, referred to as in-context training (ICT), tailors training samples to resemble few-shot prompting tasks. This prepares the model to effectively leverage a few-shot style contextual information during inference, despite the scarcity of training epitopes.

#### 4.1.1. Vanilla training

The vanilla instances (Figure 3, left) are formed as `EPI$TCR. EPI` denotes an epitope sequence, and TCR denotes a corresponding TCR sequence, sepearated by a token delimiter `$`. The model parameterized by *θ* is trained to maximize (log) likelihood of a TCR sequence *T* given a target epitope *E*, expressed as:

**Fig. 3.**
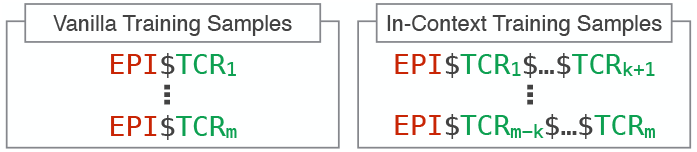
Vanilla training (Vanilla) and in-context training (ICT). The key difference lies in the context window. ICT involves multiple in-context TCRs within each training sample, allowing the model to leverage contextual TCRs when generating new sequences. However, vanilla training lacks the contextual information, relying solely on TCR-epitope pairs.

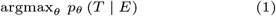

#### 4.1.2. In-context training

The ICT instances (Figure 3, right) are formed as `EPI` $`TCR` _1_$… $`TCR` _*k*+1_, leveraging multiple binding TCRs as context for each subsequent TCR. The model parameterized by *θ* is trained to maximize (log) likelihood of a TCR sequence *T*_*k*+1_ given the other TCRs *T*_1_, …, *T*_*k*_ as context and a target epitope *E*, expressed as:

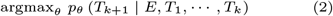

The training objective is in line with the inference objective of in-context learning via few-shot prompting (Equation 3). This refined design is intended to ultimately enhance the model’s capacity to generate high-quality TCRs for *novel epitopes* (proposed in Sections 4.2).

#### 4.1.3. Implementation detail

We fine-tune a pre-trained protein language model, `RITA m` [10], using both the vanilla and ICT training. `RITA m` consists of 300 million parameters, pre-trained on over 280 million protein sequences. For the vanilla training, we fine-tune on 96.7K unique TCR-epitope pairs, formatted as `EPI`$`TCR` (Section 3). For ICT training, we curate the samples by randomly grouping *k* + 1 TCRs that bind the same epitope into a single instance (*k* = 4 for our model), formatted as `EPI`$`TCR`_1_$… $`TCR`_*k*+1_. While the same TCR can be included in multiple instances as long as there are distinct epitopes where the same TCR is known to bind, it cannot appear multiple times with the same epitope. Comprehensive details about model training and hyperparameter tuning are provided in Section A1.1.

### 4.2. Self-contemplation prompting

Although we believe that the use of known TCRs interacting with a target epitope can lead to more reliable TCR generation, the absence of known TCRs makes in-context learning with the standard few-shot prompting (Figure 4, left) unfeasible for *novel epitopes*. To address this, we leverage a technique called self-contemplation prompting (SCP) [13]. The core idea is to have the model create its own prompts. The step-by-step process (Figure 4, right) is outlined below.

**Fig. 4.**
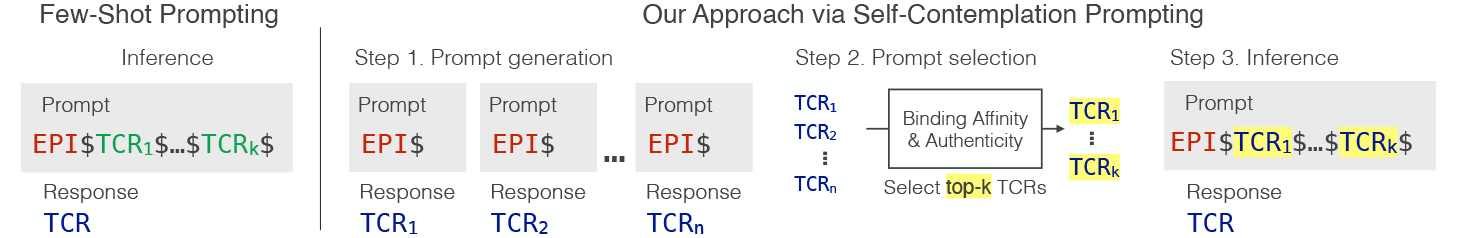
Comparison of our refined self-contemplation prompting approach (SCP) and standard few-shot prompting (FSP). Both approaches leverage TCRs interacting with a target epitope for TCR generation inference. While FSP requires known TCRs as a basis for generating new ones, SCP does not need pre-existing TCRs which makes it feasible to target novel epitopes.

#### 4.2.1. Prompt generation

First, we generate a set of *n* candidate TCRs (*n* = 300 in our experiment) for a target epitope via 0-shot prompting. The prompt simply consists of the epitope sequence followed by a delimiter (formatted as `EPI`$).

#### 4.2.2. Prompt selection

We evaluate the binding affinity and authenticity of each candidate TCR using computational models. Detailed evaluation procedure is demonstrated in Section 4.3. From this evaluation, we select best *k* high-quality TCRs to serve as prompts.

#### 4.2.3. In-context inference

The selected TCRs are combined into a *k*-shot prompt and used as input for the standard few-shot inference. The inference objective is to generate the most likley TCR sequence *t* given the other TCRs *T*_1_, …, *T*_*k*_ as context and a target epitope *E*:

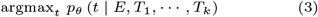

The inference objective is the same with the standard few-shot prompting (FSP) but utilizes machine-generated TCRs as a prompt, excluding the need for known TCRs.

### 4.3. TCR quality evaluation

#### 4.3.1. Metrics

We consider two key factors for evaluation of the generated TCRs: binding affinity to target epitopes and similarity to naturally occurring TCRs. These metrics act as pre-screening filters, expediting the development process by reducing the number of candidates requiring subsequent wet-lab validation.

- **Binding affinity:** Binding affinity denotes TCRs’ ability to recognize a specific antigen (target epitope). We employ two state-of-the-art computational methods to evaluate it. The first is a catELMo-based prediction model [23], referred to as *BAP*. It takes a pair of epitope and TCR sequences as input and predicts their binding affinity. The second method is *TCRMatch* [7], a pairwise scoring metric between TCRs. It provides a granular comparison of sequences on the basis of *k*-mer level similarity, where a score of 1 indicates that a pair of TCRs have the same binding profiles. Binding affinity to a target epitope is defined by the maximum similarity score of a generated TCR against ground-truth TCRs. Since TCRMatch relies on ground-truth TCRs, it is not used in the SCP prompt selection. It is only used for the final evaluation.
- **Authenticity:** Authenticity of generated TCRs, reflecting its similarity to naturally occurring TCRs, is crucial for both their functionality and safety in immunotherapy development. We use a common anomaly detection technique [18], where the likelihood of generated sequences serves as an indicator for authenticity. To this end, we fine-tune a GPT-style protein sequence model [10] to learn the patterns of naturally occurring TCRs using four million real TCR sequences from ImmunoSeq [15], named *GPT-LL*. The average log-likelihood of all amino acids within a generated TCR sequence serves as our authenticity metric.

A generated TCR is deemed high-quality, labeled ‘good’, if it exceeds predefined thresholds for both binding affinity and authenticity. To establish these optimal thresholds, we maximize Youden’s Index *J* [22]. This approach prioritizes correctly distinguishing known strong and weak binders (for TCRMatch, the binding affinity evaluation metrics) and known authentic and non-authentic sequences (for GPT-LL the authenticity evaluation metrics). The BAP threshold is further optimized to classify at least 80% of non-epitope-specific TCRs as negative, based on their distribution generated by `GPT-LL` (Section 4.3.2). These metrics serve as pre-screening filters, prioritizing high-scoring TCRs for further wet-lab functional assays. Details regarding the computational models and the process of determining optimal thresholds can be found in Section A3.

#### 4.3.2. Baselines

As no prior works exist to generate TCRs for novel epitopes, we carefully design baselines and an oracle to demonstrate unique contributions and advantages of our approach. The first baseline is a pre-trained TCR generation model trained exclusively on TCR sequences (`GPT-LL`) without epitope-specific fine-tuning, which we refer to as epitope agnostic generator. This establishes a baseline performance and helps estimate an appropriate threshold for our binding affinity metrics, under the assumption that the rate of good TCRs will be very small. The second baseline is a model fine-tuned on Vanilla training samples, performed with 0-shot inference given only the target epitope (Vanilla-0-shot). This aims to evaluate the model’s ability to generalize to unseen epitopes without any TCR information. The third and fourth baselines are models fine-tuned with ICT, and performed with few-shot inference given fake TCRs that are randomly generated from recombinations of amino acid tokens (ICT-FSP-Fake) and given randomly selected healthy TCRs known to not bind the target epitope (ICT-FSP-Healthy). Those aim to explore the impact of ICT with non-informative in-context TCRs. We also include an oracle model (ICT-FSP) that is fine-tuned with ICT and performed with few-shot inference given ground-truth TCRs. Although this scenario is not realistic for real-world applications where ground-truth is unknown, it provides a valuable estimate of the best-case few-shot performance achievable with our ICT approach.

## 5. Results

We first demonstrate that aligning training and inference through in-context training allows the model to effectively leverage contextual TCRs during inference. Furthermore, by integrating in-context training with self-contemplation prompting (SCP), we not only boost performance with contextual TCRs, but also enable the generation of high-quality TCRs without relying on ground-truth data. We also validate the reliability of our TCR evaluation metrics, ensuring robust and accurate assessments.

In our experiments, models can be exposed to data during both training and inference. For this, we classify these models based on whether each model is exposed to known binding TCRs of query epitope *E* during training and/or inference (See Table 1 for details along with section numbers each model is discussed in).

**Table 1.**
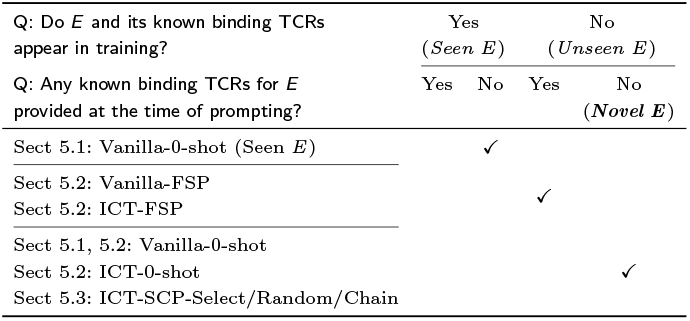
Classification of models used in all experiments based on data used in training and at the time of prompting respect to query epitope *E*.

### 5.1. TCR generation for unseen epitopes are challenging in standard framework

When there is a gap between training and inference distribution, the standard 0-shot TCR generation struggles with unseen epitopes that are never exposed to model training. This is observed across various inference temperatures (Figire 5A). At a temperature setting of 0.4, the vanilla training generates 82% of high-quality TCRs for seen epitopes it was trained on (in-sample) with a standard deviation of 0.01. However, it only generates 80% of high-quality TCRs for 29 unseen epitopes with a higher standard deviation (0.04). While the average performance between the two groups is not statistically significant, it is important to note that there are some unseen epitopes with very low success rates (Figure 5B). A similar pattern is observed for individual quality evaluation metrics such as BAP, TCRMatch and GPT-LL.

**Fig. 5.**
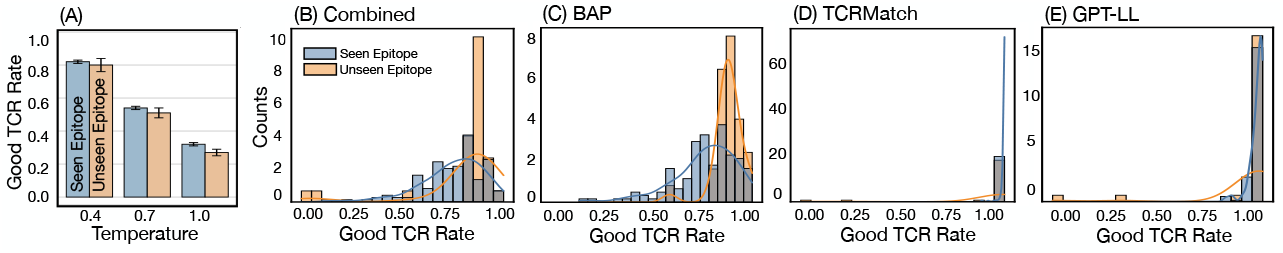
(A) The average good TCR rate of *seen* (in-sample) and *unseen* (out-of-sample) epitopes. (B–E) The good TCR rate distribution of *seen* and *unseen* epitopes (temperature 0.4).

### 5.2. In-context training is critical for few-shot TCR generation

To investigate the impact of aligning training and inference distributions, we compare vanilla training with ICT on few-shot TCR generation tasks. This approach is particularly relevant in real-world scenarios when dealing with epitopes that have very limited known binding TCRs. We hypothesize that leveraging the available in-context information during the generation process can significantly enhance TCR design by providing the model with crucial binding data as context. To isolate the effect of in-context learning, known binding TCRs for *unseen epitope* (See Table 1 for definition) are provided during inference, instead of machine generated TCRs.

We observe that vanilla training fails to leverage in-context TCRs (Figure 6). The average success rate plummets from 80% to a mere 3% when extra context TCRs are included in the prompts. Meanwhile, ICT achieves equal or better performance in k-shot than 0-shot inference for 20 out of 29 novel epitopes. Similar trends are observed with each individual metric (Figure 6). This emphasizes the crucial role of aligning the training input with how the model will be used, in order to fully unlock the potential of in-context learning for novel epitopes.

**Fig. 6.**
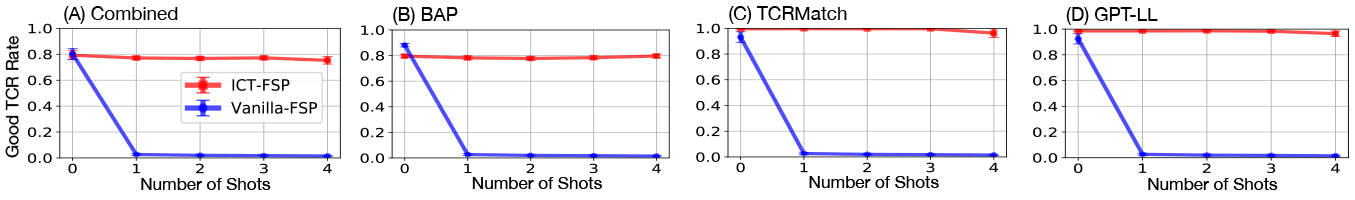
Comparison of good TCR rates across different shots for *unseen epitopes* between vanilla training with few-shot prompting (Vanilla-FSP) approach and in-context training with few-shot prompting (ICT-FSP). The key difference is in their fine-tuning samples: Vanilla training instances are formed as `EPI$TCR`, while ICT training instances are formed as EPI$TCR_1_$… $TCR_*k*+1_. Both methods use known binding TCRs as in-context samples and perform few-shot prompting during inference. These known binding TCRs are not used during training.

Visual assessment using SeqLogo also supports this observation. We select two epitopes that have shown suboptimal performance on zero-shot inference and examine their seqlogo plots generated by 4-shot inference by ICT-FSP and Vanilla-FSP against ground-truth (Figure 7). Each TCR sequence is mid-padded with a special token X, a standard method to analyze TCR sequences [12]. To evaluate differences in sequence generation patterns across models, we compared all generated sequences without filtering for “good” TCRs. This approach allows us to directly assess the intrinsic tendencies of each model’s output, providing a clearer picture of generation patterns without the influence of evaluation metrics. Vanilla training struggles to replicate the ground-truth amino acid distribution when few-shot TCR prompts are used. In contrast, TCRs generated from ICT-FSP closely mirrored the ground-truth. This disparity highlights the benefits of aligning training with inference strategies, particularly when contextual TCRs are available to guide generation.

**Fig. 7.**
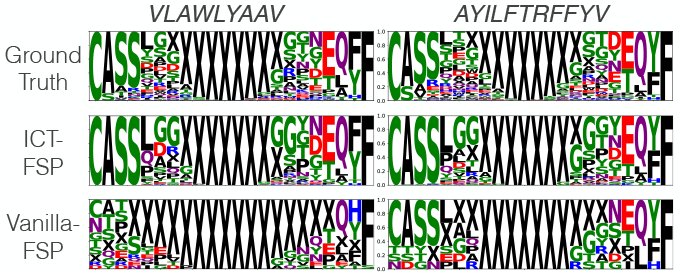
Visualization of ground-truth and generated TCRs by ICTFSP and Vanilla-FSP using a 4-shot inference using ground-truth TCRs. Each TCR sequence is mid-padded with a special token X to make the lengths of all TCRs identical prior to generating the logos. Results of the two epitopes *VLAWLYAAV* and *AYILFTRFFYV* are presented as illustration examples, selected at random.

### 5.3. Self-contemplation prompting unlock in-context TCR generation for novel epitopes

#### Carefully generated TCR prompts enhances high-quality TCR generation

By leveraging self-generated TCR sequences as in-context TCRs, our model effectively generates potential TCRs for novel epitopes without requiring known binding TCRs. We observe that SCP with longer context window (k=10) achieves a significantly higher rate of successful TCR generation compared to the baseline (Vanilla 0-shot), which lacks in-context TCRs (Table 2). We also observe that TCRs generated by SCP-Random and SCP-Select are visually similar to groundtruth TCRs (Figure A1). We also compare three SCP variants (SCP-Random, SCP-Select, and SCP-Chain). SCP-Random selects *k* in-context TCRs randomly from the model’s 0-shot inferences. SCP-Select, on the other hand, identifies high-quality TCRs and selects the top k in-context TCRs based on BAP and GPT-LL evaluation metrics. SCP-Chain iteratively selects in-context TCRs from previous inference outputs as prompts for subsequent contexts. For iteration *j*, SCP-Chain selects the best TCR generated from iteration *j* – 1 based on BAP and GPT-LL evaluation metrics, and add it to the prompt used in iteration *j* – 1. Overall, SCP-Select outperforms both SCP-Random and SCP-Chain (see Table 2). SCP-Chain often performs similarly to SCP-Random. This may be attributed to the iterative nature of SCP-Chain, where errors in earlier generations steps can propagate and accumulate.

**Table 2.**
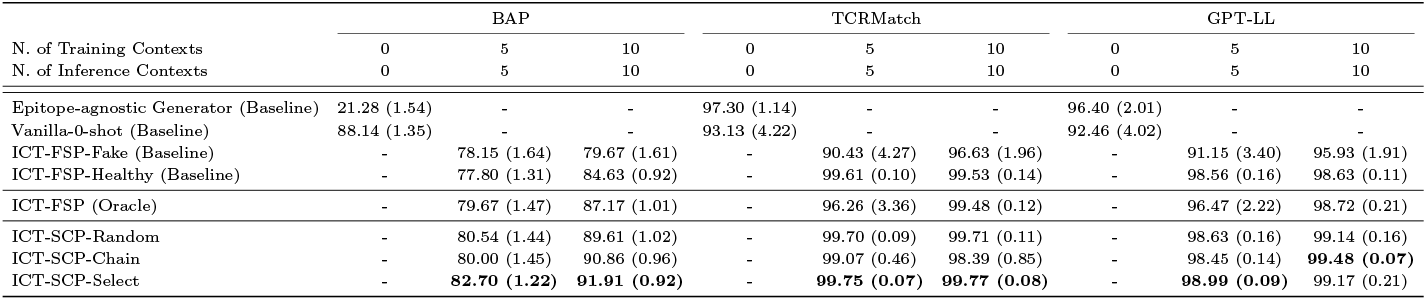
Good TCR rates of different approaches for novel epitopes. Each setting generates 300 TCRs for each epitopes.

#### Synthesized TCR can provide adequate contextual information better than or comparably to ground-truth

Given that SCP relies entirely on self-generated TCRs as in-context prompts, we compare the performance of synthesized context TCRs from SCP (ICT-SCP-Random) to ground-truth context TCRs (ICT-FSP). While ground-truth context TCRs are impractical to obtain for novel epitopes, ICT-FSP can serve as a performance upper bound. Conversely, we evaluate the performance of Fake (ICT-FSP-Fake) and healthy TCRs (ICT-FSP-Healthy) as non-binding inference contexts, providing a performance lower bound. Table 2 shows that ICT-SCP-Random performs better than or comparably to ICT-FSP, suggesting that SCP, even with self-generated TCRs, is as effective as models utilizing ground-truth binding TCRs as inference context. This ability to iteratively generate its own prompts makes SCP independent of pre-existing TCR knowledge and expands its applicability to cases lacking known binding TCRs. Our findings signify SCP’s effectiveness, particularly when combined with in-context training, for generating TCRs in scenarios where ground-truth TCR data is unavailable.

#### A minimal number of context TCRs are sufficient to achieve optimal performance

The performance of TCR generation for novel epitopes increases with the number of context TCRs provided as inference prompts and converges after a small number of shots (Figure 8). BAP reached its performance limits after six shots. However, TCRMatch and GPT-LL seems to have already reached its performance ceiling at 1-shot. This is presumably because they exhibit strong performance in discriminating non-binding sequences (Figure 9B) and the model is already capable of generating TCRs that deviate from random distributions only based on a target epitope.

**Fig. 8.**
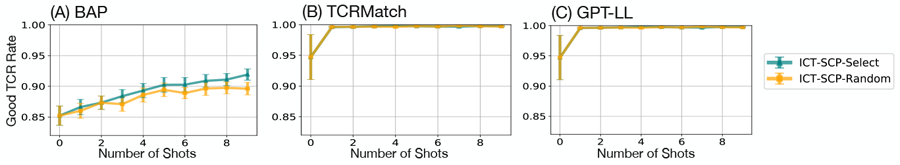
Good TCR rates of self-contemplation prompting (SCP) approaches for novel epitopes along with the number of inference shots (i.e., context TCRs). All inferences are performed by a model trained on in-context instances formed as formed as EPI$TCR_1_$… $TCR_10_. Each setting generates 300 TCRs for each epitopes.

**Fig. 9.**
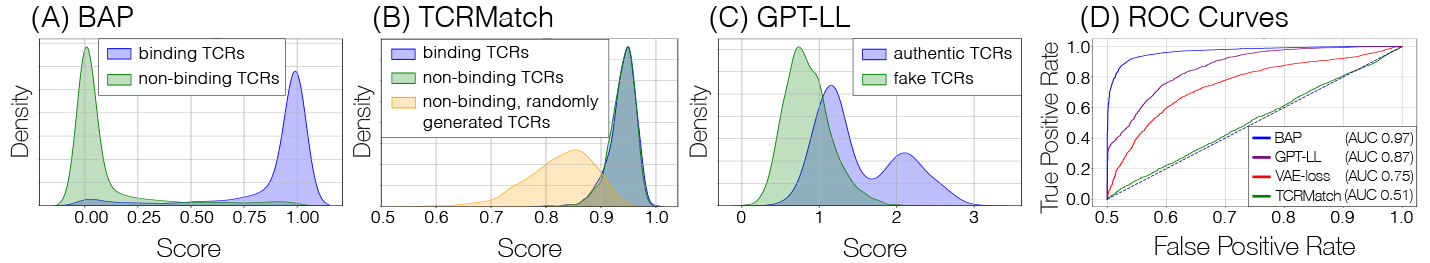
Distribution of the TCR quality evaluation metrics. (A) BAP, a measure of binding affinity, is assessed by comparing the scores of binding TCRs against non-binding healthy TCRs given a set of target epitopes. (B) TCRMatch, another measure of binding affinity, is assessed by comparing the scores of three TCR groups given a set of target epitopes: binding TCRs, non-binding healthy TCRs, and non-binding randomly generated TCRs. (C) GPT-LL, a measure of TCR authenticity, is assessed by comparing the scores (log-likelihoods) of authentic TCRs obtained from a healthy repertoire with those of randomly generated fake TCRs. (D) ROC curves illustrating the predictive power of the TCR quality evaluation metrics in distinguishing the high and low-quality TCRs.

### 5.4. Reliability of TCR quality metrics

We assess the reliability of three TCR quality evaluation metrics: BAP, TCRMatch, and GPT-LL by (1) statistically testing whether metric scores of high-quality and low-quality TCR groups originated from different distributions and (2) evaluating the ability of each metric to accurately predict the high-quality and low-quality TCRs (Figure 9). For binding affinity metrics (BAP and TCRMatch), we compare the distribution of curated binding (high-quality) and non-binding (low-quality) TCRs. Similarly, the authenticity metric (GPT-LL) was assessed using authentic (high-quality) and fake (low-quality) TCRs. Details about data curation and optimal threshold are provided in Section A3.3.

#### 5.4.1. Statistical comparison

The Kolmogorov-Smirnov (KS) test indicates that BAP scores of the binding and non-binding TCRs have different underlying distributions (Figure 9 A, p-value < 0.0001). Similarly, GPT-LL effectively discriminates between authentic and fake TCRs, with the test confirming a significant difference in their underlying distributions (Figure 9C, p-value < 0.0001). Although the difference in distributions for the TCRMatch metric is less pronounced (p-value = 0.09), it remains valuable, particularly since it significantly identifies non-binding, potentially nonsense TCR sequences such as S and AA (p-value < 0.0001, also visualized in Figure 9B).

#### 5.4.2. Predictive power

BAP has been proven to perform well in distinguishing between known binders and non-binders (AUC 0.97) (Figure 9D). GPT-LL also shows robust performance with an AUC of 0.87, effectively predicting the authenticity of TCR sequences. While TCRMatch has a lower AUC of 0.51, it is included for its proficiency in identifying non-binding sequences with random combinations such as ZYZSTS, SAVBYDCCG, and DPTZEBNSTR. Although we considered VAE-loss (detailed in Section A3) as a potential metric for authenticity, it was ultimately excluded because it did not provide additional discriminative value beyond the insights offered by the other three metrics, despite its moderate AUC of 0.75.

## 6. Discussion and Conclusion

Our study is the first to delve into high-quality TCR generation especially for novel epitopes. The challenge is, due to scarce data and limited diversity, TCR sequence generation models exhibit limited generalizability. These models fail to take advantage of in-context learning when generating binding TCRs for novel epitopes, which typically come from a different data distribution. To address this, we shape training samples into a form of few-shot learning, reducing the discrepancy between training and inference distributions. Since no known binding TCRs exist for novel epitopes, we generate and select high-quality TCRs to serve as a few-shot prompt. This ultimately improves the model’s performance in few-shot settings. We also introduce a reliable way for screening the generated TCRs based on two key criteria: binding affinity and authenticity. These screened TCRs become strong candidates for further wet-lab validation to confirm their effectiveness.

This method of designing high-quality TCRs for novel epitopes (new targets) is crucial to accelerate the design of personalized immunotherapy strategies. Computational generating and evaluating TCRs significantly speed up the T cell engineering process. This results in substantial cost reductions by minimizing the need for labor-intensive wet-lab validation.

Our findings, while specifically focused on the generation of TCR sequences targeting a particular epitope, may offer valuable insights for the broader field of biological sequence generation. The key finding is the importance of both context-rich training and inference. This includes training a model in the same way as inference and incorporating synthetic data to enrich the inference when real-world data is limited. This might be beneficial for many biological sequence generation tasks that often grapple with challenges such as limited dataset size, constrained diversity, and strong emphasis on targeted generation.

While we provides specific methods for determining cutoffs, alternative approaches, such as t-tests, could be explored. The thresholds used are illustrative and should be adjusted based on users’ tolerance for false positives. Higher thresholds reduce false positives but may miss TCRs of interest, while lower thresholds capture more TCRs at the cost of increased false positives. Further, the evaluation metrics used in this study, while achieving high precision, are not without inherent error. Ultimately, wet-lab validation remains a definitive step in confirming the TCR effectiveness. However, it is important to note that this computational pipeline prioritizes curating a refined set of high-quality candidate TCRs for subsequent wet-lab validation, resulting in expediting overall TCR engineering process.

## 7. Competing interests

The authors have no competing interests to declare that are relevant to the content of this article.

## 8. Author contributions statement

H.L. conceived the idea for the overall project. P.Z., S.B. and H.L. conceived the approach and the experiments and wrote the manuscript. P.Z. conducted the experiments.

## 9. Acknowledgments

This work was supported in part by the National Institute of General Medical Sciences of the National Institutes of Health under award number R35GM155417.

## A1 Model training

### A1.1 Design and training

Our models are fine-tuned from the `RITA_m`, a large pretrained protein language models (pLMs). It is a medium-sized model with 300 million parameters across 24 layers, selected for its optimal balance between computational efficiency and generative performance. A character-level tokenizer is used to process epitope and TCR sequences, treating each amino acid as a distinct token. Special tokens such as `$` (delimiter), `<EOS>` and `<PAD>` are included to manage sequence operations effectively within our model’s architecture. We employ a cross-entropy loss function focused on predicting tokens following the delimiter.

### A1.2 Hyperparameter tuning

We optimize the model with the following search space (bold indicates our final choices): number of contex-utual TCRs (`T`) for ICT – {3, **5**, 10 }, batch size – {**16**, 32 }, learning rate – {2 × 10^−4^, **2** × **10**^−**5**^, 1 × 10^−5^ }, training epoch – {**1**, 2, 4}, and temperature – {**0.4**, 0.7, 1.0 }. Training is stopped after the first epoch because validation error began to increase. We train the model via Adam optimizer with linear learning rate scheduler. The hyper-parameters are tuned via grid search.

### A1.3 Computing resources

The model is implemented using Pytorch with torch version 2.0.0 and CUDA version 11.7. It is trained on two NVIDIA GTX 2080 Ti GPUs in parallel (VRAM about 11GB) with training times of under 3 hours per epoch.

## A2 Model inference

During inference, the model generates TCR sequences by extending beyond the delimiter token using beam search. To control the generation process, we set a maximum sequence length of 64 amino acids, a `top_k` sampling value of 8 to encourage diversity.

## A3 TCR quality evaluation

We introduce two key aspects that are commonly used to evaluate quality of generated TCRs: binding affinity to the target epitope and similarity to naturally occurring TCRs. These metrics act as pre-screening filters, expediting the development process by reducing the number of candidates requiring subsequent wet-lab validation.

### A3.1 Binding affinity

- **BAP**: BAP [9] is a binding affinity prediction model that takes a pair of epitope and TCR sequences as input and outputs the predicted probability of binding. It employs a catELMo embedding [9] for TCR sequences and a traditional BLOSUM embedding [4] for epitope sequences. These embeddings are then fed into separate neural network layers with SiLU activation, batch normalization, and dropout. They are then concatenated and fed into another neural network with similar layers. Finally, a single neuron with a sigmoid activation function outputs a binding affinity score between 0 and 1 where a score of 0 indicates no predicted binding affinity, while 1 signifying a strong binding potential.
- **TCRMatch**: TCRMatch [1], is a *k*-mer matching algorithm that assesses the similarity between two TCR sequences by progressively breaking them down into *k*-mers (i.e., *k* amino acids). The final score is a normalized sum of similarity scores for all *k*-mer sizes. To estimate how well a generated TCR might bind to a specific epitope, we compare it to a set of ground-truth TCRs that bind to the target epitope. We first randomly select 50 of these known TCRs and calculate a TCRMatch score for the generated TCR against each of these ground-truth TCRs. The final binding affinity score is the highest among these 50 TCRMatch scores. This score ranges from 0 to 1, where 0 indicates no sequence similarity and 1 denotes a strong similarity to known binding TCRs.

### A3.2 Authenticity

- **GPT-LL**: GPT-LL is a GPT-style model fine-tuned on a protein sequence model [5] to learn patterns present in four million real TCR sequences from TCR repertoire data (ImmunoSeq, [7]). It comprises 24 transformer layers and a total of 300 million parameters, outputting a probability distribution over all possible amino acids in the vocabulary. Naturally occurring TCRs exhibit unique patterns due to their generation process: they are formed by recombining three different gene segments of multiple alleles and incorporating random base substitutions at the junctions. The log-likelihood score from a model trained on real TCRs can help identify these characteristic patterns. We calculate the GPT-LL log-likelikhood of each amino acid token in the generated TCR sequences and average these values to obtain a measure of authenticity. This score ranges from -Inf to Inf. Lower log-likelihood scores indicate a higher degree of anomaly, meaning the sequence is less likely to be a real TCR. Conversely, higher log-likelihood scores suggest a higher degree of authenticity.
- **VAE-loss**: We use a common technique using a VAE reconstruction loss as an indicator of generated output authenticity. Given a VAE’s ability to encode input data (TCR sequences in our setting) into a compressed latent space and reconstruct it back, authentic TCR sequences are expected to have low reconstruction loss. We adopt a VAE architecture proposed by [2], adapting the input format and training dataset since their model was designed for CDR3 sequences along with V and J genes, whereas we only require CDR3 sequences as input. The model utilizes BLOSUM62 matrix to embed TCR sequences and feeds them into bidirectional LSTM layers. The objective function jointly minimizes reconstruction loss (measuring the accuracy of sequence regeneration from the latent space) and Kullback–Leibler divergence (quantifying the similarity between generated and real data distributions). We optimize the model using the Adam optimizer, and implement an early stopping to prevent overfitting, halting training after 30 epochs without validation loss improvement or reaching a maximum of 200 epochs.

### A3.3 Identification of optimal threshold

We identify the optimal thresholds for both criteria by maximizing Youden’s Index (*J*) [8]. Detailed procedure is as follow.

- **Curation of binding and non-binding TCRs**: To determine the optimal threshold for the binding affinity measures, we curate sets of binding and non-binding TCRs specific to the target epitopes. For binding TCRs, we randomly source experimentally validated TCRs known to interact with the target epitopes from our test set. For non-binding TCRs, we follow a common practice [6, 3, 9], as there is a limited availability of experimentally validated non-binding TCRs. We source TCRs from healthy repertoires (ImmunoSEQ [7]) and randomly pair with target epitopes, generating non-binding TCR-epitope pairs. This resulted in 2,900 binding and 2,900 non-binding TCR-epitope pairs (100 binding and non-binding TCRs per each target epitope).
- **Curation of authentic and fake TCR**: To determine the optimal threshold for the authenticity measure, we create two datasets: a set of authentic TCRs sourced from human repertoires and a set of synthetically generated fake TCRs. To synthesize fake TCRs, we first compute the position-specific amino acid profile of four million TCR sequences from human repertoires (ImmunoSeq). For each authentic TCR, we generate a fake TCR of the same length, by sampling an amino acid for each position from the position-specific profile. This produces fake TCR sequences with realistic amino acid frequencies at each position but lacking contextual dependencies among amino acids. As a result, we obtain 2,900 authentic TCRs and 2,900 fake TCRs.
- **Identifying the best threshold via Youden’s Index J**: We identify the optimal cut-off point for each evaluation metric by maximizing Youden’s Index (*J*): *J* = sensitivity + specificity −1. This is equivalent to maximizing the true positive rate while minimizing the false positive rate. The curated binding and non-binding TCR groups serve as positive and negative sets for determining binding affinity metrics thresholds (BAP and TCRMatch). The curated authentic and fake TCR groups are used for establishing an authenticity metric threshold (GPT-LL). A simple search iterating through a set of possible values between 0 and 1 identifies the threshold that yields the maximum value. Once these thresholds are established, a generated TCR is classified as ‘high-quality’ or ‘good’ TCR only if it surpasses all three metrics’ established thresholds.

## A4 Additional Figures and Tables

**Figure A1.**
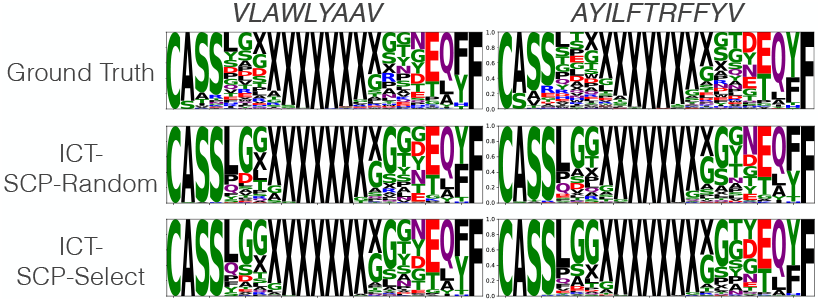
Visualization of ground-truth and generated TCRs by ICT-SCP-Random and ICT-SCP-Select using a 4-shot inference. Results of the two epitopes *VLAWLYAAV* and *AYILFTRFFYV* are presented. Notably, the SCP method operates without requiring known binding TCRs as part of the in-context prompts, relying instead on self-generated TCRs.

## Footnotes

1 Various terms have been employed to describe this technique as mentioned in Section 2. We refer to this approach as in-context training (ICT) for consistency with the established terminology of in-context learning (ICL).

2 https://github.com/Lee-CBG/TCRGen

## Notes

### Competing Interest Statement

The authors have declared no competing interest.

## References

1. Meriem Attaf, Mateusz Legut, David K Cole, and Andrew K Sewell. The T cell antigen receptor: the Swiss army knife of the immune system. Clinical & Experimental Immunology, 181(1):1–18, 2015.

2. Estelle Baulu, Ćelia Gardet, Nicolas Chuvin, and Stéphane Depil. TCR-engineered T cell therapy in solid tumors: State of the art and perspectives. Science Advances, 9(7):eadf3700, 2023.

3. Tom Brown, Benjamin Mann, Nick Ryder, Melanie Subbiah, Jared D Kaplan, Prafulla Dhariwal, Arvind Neelakantan, Pranav Shyam, Girish Sastry, Amanda Askell, et al. Language models are few-shot learners. Advances in Neural Information Processing Systems, 33:1877–1901, 2020.

4. Michael Cai, Seojin Bang, Pengfei Zhang, and Heewook Lee. ATM-TCR: TCR-epitope binding affinity prediction using a multi-head self-attention model. Frontiers in Immunology, 13, 2022.

5. Yanda Chen, Ruiqi Zhong, Sheng Zha, George Karypis, and He He. Meta-learning via language model in-context tuning. In Proceedings of the 60th Annual Meeting of the Association for Computational Linguistics (Volume 1: Long Papers), pages 719–730. Association for Computational Linguistics, 2022.

6. Ziqi Chen, Martin Renqiang Min, Hongyu Guo, Chao Cheng, Trevor Clancy, and Xia Ning. T-cell receptor optimization with reinforcement learning and mutation polices for precision immunotherapy. In International Conference on Research in Computational Molecular Biology, pages 174–191. Springer, 2023.

7. William D Chronister, Austin Crinklaw, Swapnil Mahajan, Randi Vita, Zeynep Koşaloğlu-Yalcin, Zhen Yan, Jason A Greenbaum, Leon E Jessen, Morten Nielsen, Scott Christley, et al. TCRMatch: predicting T-cell receptor specificity based on sequence similarity to previously characterized receptors. Frontiers in Immunology, 12:640725, 2021.

8. Kristian Davidsen, Branden J Olson, William S DeWitt III, Jean Feng, Elias Harkins, Philip Bradley, and Frederick A Matsen IV. Deep generative models for T cell receptor protein sequences. eLife, 8:e46935, 2019.

9. Noelia Ferruz, Steffen Schmidt, and Birte Höcker. ProtGPT2 is a deep unsupervised language model for protein design. Nature Communications, 13(1):4348, 2022.

10. Daniel Hesslow, Niccoló Zanichelli, Pascal Notin, Iacopo Poli, and Debora Marks. RITA: a study on scaling up generative protein sequence models. arXiv preprint 2205.05789, 2022.

11. Vanessa Isabell Jurtz, Leon Eyrich Jessen, Amalie Kai Bentzen, Martin Closter Jespersen, Swapnil Mahajan, Randi Vita, Kamilla Kjærgaard Jensen, Paolo Marcatili, Sine Reker Hadrup, Bjoern Peters, et al. NetTCR: sequence-based prediction of TCR binding to peptide-MHC complexes using convolutional neural networks. BioRxiv, page 433706, 2018.

12. Marie-Paule Lefranc, Christelle Pommié, Quentin Kaas, Elodie Duprat, Nathalie Bosc, Delphine Guiraudou, Christelle Jean, Manuel Ruiz, Isabelle D. Piédade, Mathieu Rouard, et al. IMGT unique numbering for immunoglobulin and T cell receptor constant domains and Ig superfamily C-like domains. Developmental & Comparative Immunology, 29(3):185–203, 2005.

13. Rui Li, Guoyin Wang, and Jiwei Li. Are human-generated demonstrations necessary for in-context learning? International Conference on Learning Representations, 2024.

14. Sewon Min, Mike Lewis, Luke Zettlemoyer, and Hannaneh Hajishirzi. MetaICL: Learning to learn in context. In Marine Carpuat, Marie-Catherine de Marneffe, and Ivan Vladimir Meza Ruiz, editors, Proceedings of the 2022 Conference of the North American Chapter of the Association for Computational Linguistics: Human Language Technologies, pages 2791–2809. Association for Computational Linguistics, July 2022.

15. Sean Nolan, Marissa Vignali, Mark Klinger, Jennifer N Dines, Ian M Kaplan, Emily Svejnoha, Tracy Craft, Katie Boland, Mitch Pesesky, Rachel M Gittelman, et al. A large-scale database of T-cell receptor beta (TCRβ) sequences and binding associations from natural and synthetic exposure to SARS-CoV-2. Research Square, 2020.

16. Mikhail Shugay, Dmitriy V Bagaev, Ivan V Zvyagin, Renske M Vroomans, Jeremy Chase Crawford, Garry Dolton, Ekaterina A Komech, Anastasiya L Sycheva, Anna E Koneva, Evgeniy S Egorov, et al. VDJdb: a curated database of T-cell receptor sequences with known antigen specificity. Nucleic Acids Research, 46(D1):D419–D427, 2018.

17. Ido Springer, Nili Tickotsky, and Yoram Louzoun. Contribution of T cell receptor alpha and beta CDR3, MHC typing, V and J genes to peptide binding prediction. Frontiers in Immunology, 12:664514, 2021.

18. Jing Su, Chufeng Jiang, Xin Jin, Yuxin Qiao, Tingsong Xiao, Hongda Ma, Rong Wei, Zhi Jing, Jiajun Xu, and Junhong Lin. Large language models for forecasting and anomaly detection: A systematic literature review. arXiv preprint 2402.10350, 2024.

19. Nili Tickotsky, Tal Sagiv, Jaime Prilusky, Eric Shifrut, and Nir Friedman. McPAS-TCR: a manually curated catalogue of pathology-associated T cell receptor sequences. Bioinformatics, 33(18):2924–2929, 2017.

20. Randi Vita, Swapnil Mahajan, James A Overton, Sandeep Kumar Dhanda, Sheridan Martini, Jason R Cantrell, Daniel K Wheeler, Alessandro Sette, and Bjoern Peters. The immune epitope database (IEDB): 2018 update. Nucleic Acids Research, 47(D1):D339–D343, 2019.

21. Muzamil Y Want, Zeenat Bashir, and Rauf A Najar. T cell based immunotherapy for cancer: approaches and strategies. Vaccines, 11(4):835, 2023.

22. William J Youden. Index for rating diagnostic tests. Cancer, 3(1):32–35, 1950.

23. Pengfei Zhang, Seojin Bang, Michael Cai, and Heewook Lee. Context-aware amino acid embedding advances analysis of TCR-epitope interactions. eLife, April 2023.

24. Pengfei Zhang, Seojin Bang, and Heewook Lee. PiTE: TCR-epitope binding affinity prediction pipeline using Transformer-based sequence encoder. In Pacific Symposium on Biocomputing 2023: Kohala Coast, Hawaii, USA, 3–7 January 2023, pages 347–358. World Scientific, 2022.

25. Pengfei Zhang, Seojin Bang, and Heewook Lee. Active learning framework for cost-effective TCR-epitope binding affinity prediction. In 2023 IEEE International Conference on Bioinformatics and Biomedicine (BIBM), pages 988–993. IEEE, 2023.

26. Zihao Zhao, Eric Wallace, Shi Feng, Dan Klein, and Sameer Singh. Calibrate before use: Improving few-shot performance of language models. In International Conference on Machine Learning, pages 12697–12706. PMLR, 2021.

## Refereneces

[1] William D Chronister, Austin Crinklaw, Swapnil Mahajan, Randi Vita, Zeynep Koşaloğlu-Yalçin, Zhen Yan, Jason A Greenbaum, Leon E Jessen, Morten Nielsen, Scott Christley, et al. TCRMatch: predicting T-cell receptor specificity based on sequence similarity to previously characterized receptors. Frontiers in Immunology, 12:640725, 2021.

[2] Kristian Davidsen, Branden J Olson, William S DeWitt III, Jean Feng, Elias Harkins, Philip Bradley, and Frederick A Matsen IV. Deep generative models for T cell receptor protein sequences. eLife, 8:e46935, 2019.

[3] Sofie Gielis, Pieter Moris, Wout Bittremieux, Nicolas De Neuter, Benson Ogunjimi, Kris Laukens, and Pieter Meysman. Detection of enriched T cell epitope specificity in full T cell receptor sequence repertoires. Frontiers in Immunology, 10:2820, 2019.

[4] Steven Henikoff and Jorja G Henikoff. Amino acid substitution matrices from protein blocks. Proceedings of the National Academy of Sciences, 89(22):10915–10919, 1992.

[5] Daniel Hesslow, Niccoló Zanichelli, Pascal Notin, Iacopo Poli, and Debora Marks. RITA: a study on scaling up generative protein sequence models. arXiv preprint 2205.05789, 2022.

[6] Vanessa Isabell Jurtz, Leon Eyrich Jessen, Amalie Kai Bentzen, Martin Closter Jespersen, Swapnil Mahajan, Randi Vita, Kamilla Kjærgaard Jensen, Paolo Marcatili, Sine Reker Hadrup, Bjoern Peters, et al. NetTCR: sequence-based prediction of TCR binding to peptide-MHC complexes using convolutional neural networks. BioRxiv, page 433706, 2018.

[7] Sean Nolan, Marissa Vignali, Mark Klinger, Jennifer N Dines, Ian M Kaplan, Emily Svejnoha, Tracy Craft, Katie Boland, Mitch Pesesky, Rachel M Gittelman, et al. A large-scale database of T-cell receptor beta (TCRβ) sequences and binding associations from natural and synthetic exposure to SARS-CoV-2. Research Square, 2020.

[8] William J Youden. Index for rating diagnostic tests. Cancer, 3(1):32–35, 1950.

[9] Pengfei Zhang, Seojin Bang, Michael Cai, and Heewook Lee. Context-aware amino acid embedding advances analysis of TCR-epitope interactions. eLife, April 2023.

